# *MolLM*: A Unified Language Model for Integrating Biomedical Text with 2D and 3D Molecular Representations

**DOI:** 10.1101/2023.11.25.568656

**Authors:** Xiangru Tang, Andrew Tran, Jeffrey Tan, Mark B. Gerstein

**Affiliations:** Yale University, New Haven, CT 06520, USA

## Abstract

**Motivation:** The current paradigm of deep learning models for the joint representation of molecules and text primarily relies on 1D or 2D molecular formats, neglecting significant 3D structural information that offers valuable physical insight. This narrow focus inhibits the models’ versatility and adaptability across a wide range of modalities. Conversely, the limited research focusing on explicit 3D representation tends to overlook textual data within the biomedical domain.

**Results:** We present a unified pre-trained language model, MolLM, that concurrently captures 2D and 3D molecular information alongside biomedical text. MolLM consists of a text Transformer encoder and a molecular Transformer encoder, designed to encode both 2D and 3D molecular structures. To support MolLM’s self-supervised pre-training, we constructed 160K molecule-text pairings. Employing contrastive learning as a supervisory signal for cross-modal information learning, MolLM demonstrates robust molecular representation capabilities across 4 downstream tasks, including cross-modality molecule and text matching, property prediction, captioning, and text-prompted molecular editing. Through ablation, we demonstrate that the inclusion of explicit 3D representations improves performance in these downstream tasks.

**Availability and implementation:** Our code, data, and pre-trained model weights are all available at https://github.com/gersteinlab/MolLM.

## 1 Introduction

Pre-training techniques in natural language processing (NLP) have resulted in significant shifts in numerous domains by enabling the interpretation of complex patterns from large-scale datasets (Radford *et al*., 2018; Vaswani *et al*., 2017; Devlin *et al*., 2019; Raffel *et al*., 2020; Wei *et al*., 2021). This success has led to promising outcomes in the biomedical field, particularly in tasks such as molecular property prediction and chemical structure generation, indicating a strong artificial intelligence potential in biomolecular studies (Gu *et al*., 2021; Xia *et al*., 2022; Luo *et al*., 2022a; Singhal *et al*., 2023).

Past endeavors have sought to learn representations of molecular chemical properties through supervised tasks using labeled data (Hwang *et al*., 2020; Zuranski *et al*., 2021). Nevertheless, these efforts heavily rely on the availability of large-scale datasets, prompting an increased focus on self-supervised learning. Recent investigations have leveraged pre-trained, self-supervised methods to derive representations of molecular structures (Zeng *et al*., 2022; Rong *et al*., 2020; Zhang *et al*., 2021), primarily employing a masking approach to learning the geometric structure and consequently obtaining corresponding chemical structure representations. However, these endeavors have predominantly concentrated on 2D molecular structures (Su *et al*., 2022; Pogány *et al*., 2018; Coley *et al*., 2017; Liu *et al*., 2023a), thereby neglecting the pivotal role of 3D structures in molecular modeling. In practical terms, 3D molecular structures encapsulate crucial chemical information, and the positions of functional groups within them serve as potent predictors for understanding molecular properties and modeling interactions. While some research has incorporated 3D geometric structures, these works have their limitations in model architecture (Yu *et al*., 2021; Miao *et al*., 2023; Jiang *et al*., 2021; Devinyak *et al*., 2014).

Additionally, the natural language modality encompasses a vast volume of biochemical knowledge, offering a valuable resource for enhancing model comprehension of molecular structures. When applied to downstream tasks, a model’s proficiency in interpreting natural language facilitates more refined control of molecule generation and editing, using natural language as the mode of interaction. Notably, recent state-of-the-art multimodal models for molecular tasks do not incorporate essential 3D information alongside natural language (Su *et al*., 2022). Hence, this underscores the imperative to develop models that can seamlessly integrate both 2D and 3D molecular structures with biomedical text to propel scientific discovery. Inspired by this, we introduce MolLM, the first molecular multimodal language model that combines 2D and 3D molecular structures with natural language—a novel approach that has not been explored before.

In this work, we first create a multimodal dataset for self-supervised pre-training, including molecules sourced from PubChem (Kim *et al*., 2023) and related publicly available academic texts, totaling 160K molecule graph-text pairs. At the core of MolLM is a unified pre-training framework that seamlessly blends a text Transformer encoder with a molecular Transformer encoder, pre-trained on molecular graphs and related textual data jointly. Specifically, the text Transformer encoder is pre-trained on biomedical text. To effectively capture the multimodal molecular structure information, we then adopt a molecular graph Transformer encoder. Leveraging the graph Transformer architecture, we integrate pair-wise encodings to represent both 2D graphs and 3D geometric structures. In other words, we integrate positional encoding with the standard graph encoding, allowing the model to capture atom-wise structural information. Following Transformer-M (Luo *et al*., 2022b), we effectively leverage both 2D and 3D molecular structures for better performance in downstream tasks. Crucially, our model aligns these cross-modal representations during pre-training through the paradigm of contrastive learning. Contrastive learning efficiently facilitates self-supervised pre-training by bringing similar molecules closer together and placing dissimilar molecules farther apart in the representation space.

Our presented work further emphasizes the significance of capturing 3D molecular structures. MolLM overcomes the limitations of existing pre-trained molecular models, which can only process different data types separately, by offering a comprehensive understanding of molecular entities. We have conducted extensive experiments to showcase the model’s capacity to capture molecular expertise, demonstrating promising performance in downstream tasks such as (1) **cross-modal retrieval** (retrieving the related molecule or text with the query of the opposite modality), (2) **molecule captioning** (generating natural language descriptions of molecules), (3) **property prediction** (classifying molecule properties), and (4) **molecule editing** (editing molecules based on natural language prompts) (Zeng *et al*., 2022; Edwards *et al*., 2022; Wu *et al*., 2017a; Liu *et al*., 2023b). Our promising results demonstrate that integrating 3D information is beneficial. For example, in the cross-modality matching task, while our multimodal architecture outperforms all baselines in various zero-shot and supervised settings, the direct incorporation of 3D data further improves each score. In summary, to the best of our knowledge, this is the first work that introduces a 2D and 3D pre-training framework for molecules in conjunction with textual information, presenting the following two significant contributions to the field. Firstly, we introduce a novel dataset, surpassing baseline datasets in size while maintaining comparable text complexity. Our results underscore the necessity of datasets containing profound textual information and substantial scale for maximizing effectiveness. Secondly, we demonstrate the efficacy of multimodal pre-trained models in molecular analysis, particularly when incorporating all three modalities of textual, 2D, and 3D information.

## 2 Related work

### 2.1 Molecular Representations

Historically, many works have focused on 1D molecular representations, with the Simplified Molecular Input Line Entry System (SMILES) being the most common method (Weininger, 1988). Many papers have leveraged SMILES for a variety of molecular tasks (Chen and Zhang, 2021; Kuenzi *et al*., 2020; An *et al*., 2022). For a more explicit representation of molecules, graph-based representations have become a popular method for integrating 2D structures. The most common and straightforward graph representation involves vertices representing atoms and their respective features and edges representing bonds and their respective features. Utilizing Open Graph Benchmark (Hu *et al*., 2020), various works have applied common graph model architectures for molecule analysis, such as CNNs, graph convolutional networks (GCNs), and GraphSAGE. (Pogány *et al*., 2018; Coley *et al*., 2017; Liu *et al*., 2023a). While these models achieve good results, they fail to account for nuances within the 3D molecular structure, such as stereochemistry, interaction energies, and spatial binding affinities. More recent works include 3D molecular graph data by providing 3D coordinates of each molecule within the atom representation. As 3D data are not universally available for many molecules, most studies generate 3D conformations using molecular analysis libraries such as RDKit (Landrum, 2023). By focusing on 3D contexts, these models demonstrate improved performance on certain tasks where 3D structure significantly impacts outcomes, such as property prediction (Kuzminykh *et al*., 2018; Thomas *et al*., 2018; Kajita *et al*., 2017; Stärk *et al*., 2022; Li *et al*., 2022b; Yang *et al*., 2019). Some models use contrastive learning techniques to accommodate multiple modalities of molecule inputs, aiming to unify 2D and 3D structural information (Liu *et al*., 2022; Zhu *et al*., 2022; Li *et al*., 2022a).

### 2.2 Molecular Pre-trained Language Models

The popularity of pre-trained language models has led to prior utilization of these models on molecules represented via textual formats such as SMILES. One of the initial models to explore inputting SMILES strings into language models is SMILES-BERT (Wang *et al*., 2019), which trained on masking tasks to predict missing tokens in sequences representing text, including SMILES strings alongside the regular text. Similarly, BARTSmiles expanded upon this approach by using a BART-like model and incorporating more downstream tasks (Chilingaryan *et al*., 2022). However, as these models use inherently 1D textual representations of molecules, they lose a significant amount of information regarding geometric and stereoelectric effects.

In terms of multimodality works, prior research has focused on multimodal text and molecule models to represent molecules and their related textual descriptions in a joint latent space. KV-PLM (Zeng *et al*., 2022) utilizes the SMILES representation of molecules before projecting them onto such a joint latent space, while MoMu (Su *et al*., 2022) utilizes a graph representation of molecules. These methods, along with other recent works (Wang *et al*., 2022a,b), utilize a contrastive loss function to align molecular representations and textual representations. To improve accuracy and robustness during training, MoMu employs augmentations of the molecular graphs to create semantically similar graphs (Liu *et al*., 2023b). However, due to both MoMu and KV-PLM’s molecular representation inputs, these models only consider a limited 2D view of molecular geometry.

Regarding training data, KV-PLM and MoMu each utilized datasets of 15K molecule-text pairings. KV-PLM sourced the textual data from PubChem, while MoMu sourced a different set of 15K pairs from the S2ORC (Lo *et al*., 2020) corpus dataset. As the S2ORC dataset lacks clear pairing between molecules and related text, Su *et al*. expressed concerns that simply selecting texts referencing molecules might not guarantee strongly related text. Therefore, limitations in the datasets of prior works extend beyond concerns about their size, as these datasets may also exhibit weak correlations within their molecule and text pairings.

### 2.3 Models for Specific Molecular Tasks

Molecular property prediction is one of the most widely researched molecular tasks due to its potential to expedite traditionally expensive and time-consuming wet-lab experiments. Previous works have aimed to develop models that perform well in this prediction task (Wang *et al*., 2022b; Liu *et al*., 2023c; Wen *et al*., 2022; Ross *et al*., 2022). For example, MolCLR (Wang *et al*., 2022b) performs contrastive learning via graph neural networks (GNNs) and data augmentation, however, it is not annotated with any text descriptions, limiting MolCLR’s ability to perform molecular tasks involving text.

Concerning other models capable of downstream property prediction, the representation learning method described in (Zhu *et al*., 2022) combines 2D and 3D graph data. This method encodes atomic coordinates and interatomic distances, subsequently merging them with atom representations through GNNs. Demonstrating a significant improvement over 2D-only methods on downstream property prediction tasks, this method underscores the importance of 3D representations and proposes an approach to incorporate 3D information into the pre-training process. However, as this model solely operates with molecule data without incorporating textual data as input, it lacks flexibility for more generalized tasks, many of which involve textual data.

Although molecular property prediction does not require the use of text, several works have proposed molecular tasks that necessitate a combined understanding of both molecules and text. For instance, MolT5 introduced the molecule captioning task, which involves generating meaningful textual descriptions of molecules, capturing their most important properties, structural elements, and applications (Edwards *et al*., 2022). Furthermore, MoleculeSTM introduced a molecule editing task, wherein the input comprises a molecule and a prompt, such as “this molecule is more soluble,” aiming for an output of a molecule similar to the original but aligned with the prompt (Liu *et al*., 2023b).

## 3 Methods

### 3.1 Data

To facilitate self-supervised pre-training of molecules, a large-scale text-molecule paired dataset was necessary. Consequently, we collected our dataset from two sources. First, for collecting the molecular data and directly related descriptions, we accessed the PubChem molecular database (Kim *et al*., 2023), which hosts information on over 100 million molecular compounds. From this repository, we used the PubChem Classification Browser to identify 160K molecules with substantial physical descriptions, emphasizing the comprehensiveness of the “Description” field. For example, we utilized the PubChem Classification Browser to acquire molecules such as “acetylcarnitine.” Then, using the PubChem REST API, we queried for 2D graphical data, as well as the basic physical descriptions available from PubChem.

To collect associated textual data, we followed the matching methodology employed by MoMu to extract sentences from papers within the S2ORC corpus database (Lo *et al*., 2020) relevant to certain molecules. We then paired these text descriptions with the 2D and 3D graphical representations of each molecule to create molecular graph-text pairs. Specifically, using the aforementioned example, we looked into the S2ORC database to find papers related to “acetylcarnitine,” extracting text containing the molecule, or its synonyms, from the Abstract, Introduction, and Conclusion sections of each paper. Due to the length limit of the text encoder, we constructed our textual data with a cut-off limit of 256 tokens.

Notably, while MoleculeSTM (Liu *et al*., 2023b) has a large dataset with both 2D and 3D data, it does not incorporate academic textual data, thus overlooking a significant data source. Table 1 provides a comparison of our dataset against similar datasets. Our dataset is the largest within this domain, encompassing academic text and comprehensive multimodal molecule information.

**Table 1.**
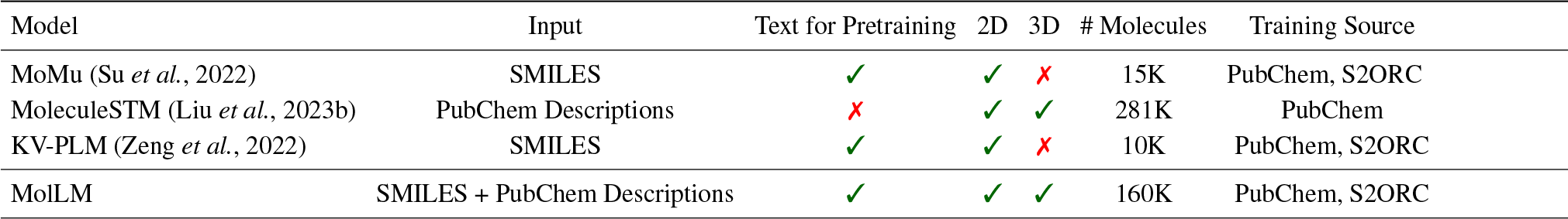
Comparison of datasets used in our study (MolLM) and the baseline approaches in this field. Each dataset is characterized by (columns 2-7) input data type, availability of text pre-training, availability of 2D/3D structural information pre-training, total molecule count, and the pre-trained data source.

| Model | Input | Text for Pretraining | 2D | 3D | # Molecules | Training Source |
| --- | --- | --- | --- | --- | --- | --- |
| MoMu (Su et al., 2022) | SMILES | ✓ | ✓ | ✗ | 15K | PubChem, S2ORC |
| MoleculeSTM (Liu et al., 2023b) | PubChem Descriptions | ✗ | ✓ | ✓ | 281K | PubChem |
| KV-PLM (Zeng et al., 2022) | SMILES | ✓ | ✓ | ✗ | 10K | PubChem, S2ORC |
| MolLM | SMILES + PubChem Descriptions | ✓ | ✓ | ✓ | 160K | PubChem, S2ORC |

### 3.2 Architecture

To support encoding both molecules and text into MolLM’s joint latent space, we have a text encoder for the textual descriptions of molecules and separately a molecule encoder for the 2D/3D representations of molecules as shown in Figure 1.

**Fig. 1.**
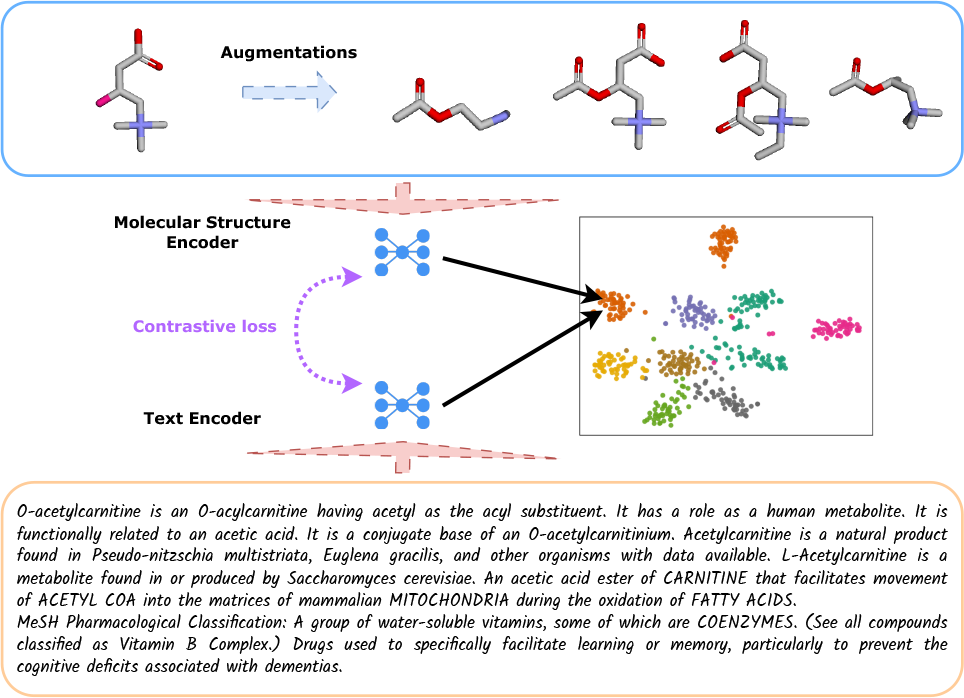
Overview of the pre-training pipeline. MolLM encodes both molecular graph data (top) and textual data (bottom). For each molecule, we generate up to four molecule augmentations before encoding the 2D and 3D structural data. On the text side, we encode related sentences from scientific journals. These encoded representations are then used during the pre-training for contrastive learning.

#### 3.2.1 Text Encoder

We initialize our text encoder’s model weights with KV-PLM’s checkpoint (Zeng *et al*., 2022), aiming to generate embeddings for the extracted sentences from academic literature in our dataset. KV-PLM enhances BERT (Bidirectional Encoder Representations from Transformers) (Devlin *et al*., 2019), a widely used general text encoder, by fine-tuning its performance on academic texts related to molecules that include SMILES strings (Weininger, 1988).

We also double the tokenizer output length limit within KV-PLM’s use of the BERT tokenizer, increasing it from 128 tokens to 256 tokens. This adjustment helps avoid unnecessary truncation and loss of detail when processing text sequences, allowing for the inclusion of more detailed text descriptions. Thus, our approach enables efficient capture of the core meaning and context of the full sentences and paragraphs.

#### 3.2.2 Molecular Structure Encoder

MolLM is designed to obtain representations of molecular data via pathways that process both 2D and 3D structural information. The 2D pathway harnesses information extracted from the molecule’s 2D graph structure, including degrees, shortest path distances, and edges. These data elements represent the spatial relationships between atoms in the molecule, enabling the model to understand its 2D architecture. On the other hand, the 3D pathway focuses on the molecule’s 3D geometrical structure, computing spatial distances among atoms within the 3D structure. This geometric perspective empowers the model to gain insights into the 3D arrangement of atoms and their complex interactions, which are often crucial in predicting molecular properties and interactions. Together, these three pathways equip MolLM with a thorough comprehension of the molecule, spanning its sequence and both its 2D and 3D structures, thereby producing a rich, multimodal representation.

Inspired by Transformer-M (Luo *et al*., 2022b), MolLM employs a similar strategy to process both 2D graph structures and 3D geometric structures in molecular representations within a unified model. Using a graph Transformer as its base, MolLM encodes structural data such as edge features, bond type, and 3D spatial relation as bias values within the attention mechanism. The Transformer uses these values to weigh and combine different parts of the input. We implement this modification in the attention mechanism because it allows the relationship between atoms in the model to be directly influenced by their 2D and 3D spatial relations, aligning with the physical significance of these geometric relations.

To summarize the implementation, we recap the encodings of these structural features that we adopt from Transformer-M. In terms of the graph structure, the graph has vertices representing atoms and edges representing the bonds between these atoms.

For encoding edge features, we utilize the 2D graph structure by considering edge feature row vectors, **e**_1_, **e**_2_, …, along the shortest path between any two vertices. These edge feature vectors represent characteristics of the chemical bonds, including bond type and stereochemistry. Suppose the edge feature vectors are of dimension *k*_*edge*_, as in *e*_*i*_ ∈ ℝ*k*^*edge*^ for each edge. Then, the element-wise average of them is denoted as *avg*(**e**). We also utilize a learnable vector **w** ∈ ℝ*k*^*edge*^ to emphasize the importance of certain bond features over others. Our encoding of edge features for any two vertices indexed as *i* and *j*, which is a scalar quantity, is given as

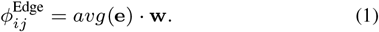

Additionally, we aim to include 2D distance, specifically the shortest path between any two atoms by following their bonds, in our structural encoding. Thus, for the relationship between atoms in the 2D space, we define an encoding to represent the shortest path distance between two atoms. Moreover, we introduce a learnable scalar *w* to allow for the weighing of the importance of considering shortest path distances in our structural encoding. We define *spd*(*i, j*) as the shortest path distance between atoms *i* and *j* in the 2D graph, counted by edges, making it a scalar quantity. Hence, our encoding for shortest path distances, also a scalar quantity, is

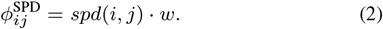

Similarly, we consider 3D spatial relations. Instead of utilizing standard Euclidean distance, we employ the Gaussian Basis Kernel as it provides a non-linear function of distance, offering a more nuanced representation of the 3D spatial relationship. Suppose we utilize *K* Gaussian Basis Kernels to represent 3D distance with the vector *ψ*(**i, j**) = [*ψ*(*i, j*, 1), …, *ψ*(*i, j, K*)] ∈ ℝ*k*. We also introduce a learnable matrix **W**_1_ ∈ ℝ^*K×K*^ and a vector **w**_2_ ∈ ℝ*k* for a multi-layer perceptron (MLP) to obtain the final encoding by processing the distance representation given by the Gaussian Basis Kernel. The MLP allows for a richer representation with higher-level features that can be inferred from the 3D distance along with the ability to selectively emphasize certain aspects, such as specific directions within the 3D distance. We utilize ReLU as the activation function, *act*, within this MLP. The 3D distance encoding is processed between vertices, indexed as *i* and *j*, as a scalar quantity, after applying the Gaussian Basis Kernel as

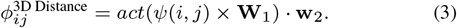

Specifically, the Gaussian Basis Kernel has a hyperparameter *σ* that controls the width of the kernel, which adjusts the sensitivity of the kernel to distance and the smoothness of its output. These aspects of the kernel align with the scales of distances at the molecular level and the potential noise of these distances, respectively. The *k*-th Gaussian Basis Kernel (Scholkopf *et al*., 1997) within *ψ*(*i, j*) is given as

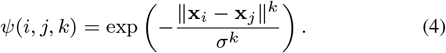

These 2D and 3D positional encodings from the graph structure of the molecule and the 3D geometric structures are linearly combined to consider them in aggregate. We incorporate this combined encoding into the attention mechanism of the graph Transformer layers. This linear combination, considering all atoms in the molecular graph, is given as

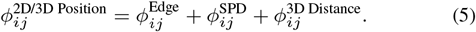

Let *a* be the atom count in the molecular graph. We define the matrix *ϕ*^2D/3D Position^ ∈ ℝ^*a×a*^ by Equation 5. We directly add this matrix as a bias to the standard Transformer attention calculation before the final application of Softmax. The motivation behind adding this matrix to the Transformer attention mechanism is to capture contextual relationships among our graph’s vertices and atoms while weighing the importance of each in the final embedding. By incorporating our bias, which is based upon 2D and 3D spatial data, the model directly considers this rich spatial data in the attention calculation, thus considering spatial relationships between atoms within the molecular structure.

### 3.3. Pre-training

For MolLM’s pretraining, data augmentation strategies allow us to enhance robustness and efficiency in handling molecular-related tasks, which has been demonstrated in (Su *et al*., 2022). We utilize contrastive learning as the objective which aims to derive meaningful representations by comparing pairs of positive and negative samples. Our approach involves two types of losses: (1) *a contrastive loss* which compares representations of different modalities within the same data sample, and (2) *a self loss* which contrasts different augmentations of the same modality within the same data sample. We define these two losses in section 3.3.2.

#### 3.3.1 Data Augmentation

We expand upon the data augmentations proposed in MoMu (Su *et al*., 2022) by introducing two augmentations that alter each molecule’s chemical features, such as molecular motifs and functional groups. These additions complement the two augmentations used in MoMu, which alter graph features like nodes and edges. In total, we apply four data augmentations to each 2D molecular graph: *(1) node dropping, (2) subgraph sampling, (3) chemical transformation*, and *(4) substructure removal*. We then compute and insert 3D atomic positions into these augmented graphs, aiming to construct semantically consistent molecular structures suitable for pre-training.

*Node dropping* randomly discards vertices from the original graph, while *subgraph sampling* involves traversing random walks. *Chemical transformation* augmentations generate new structures by applying various chemical transformations alongside other graph augmentations. These transformations entail adding and removing functional groups, followed by the optimization of 3D structures. *Substructure removal* uses BRICS decomposition to remove certain substructures in a molecular viable manner. From the possible substructure generations, a random viable molecular motif is chosen. Fig 1 provides a visualization of the augmentations and Appendix 3 shows more examples. Even with some augmentations modifying the chemical structure and properties, we consider the new instances to belong to the same label. Recent work has demonstrated the effectiveness of such augmentations during pre-training, improving model robustness, generalization, and understanding of functional groups and chemical structure compositions.

For the aforementioned augmentations, we perform manipulations and optimizations on molecular structures using the open-source RDKit library. These augmentations involve a deliberate randomization process and are repeated until we obtain an augmented molecule that measures a difference below 3.0 root-mean-square deviation (RMSD) according to RDKit’s molecular distance, or inverse similarity, calculation. We chose this threshold of 3.0 based on expert annotation and discussions with two chemistry professors, after examining the similarity of 100 augmented molecules for various RMSD values. This criterion helps avoid augmenting molecules to be too semantically different, with too large of a change of molecular properties, from the original. See Appendix 2 for further motivation behind our use of RDKit. Furthermore, for each molecule within the training dataset, only the four augmented versions are input into training while the original molecule is not. There may be a concern with not using the original molecule but our proportion of atoms to drop for *node dropping* is a conservative 10%.

Given a mini-batch of *N* molecular graphs, we generate four different augmentations for each graph. Let 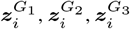, and 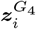 each denote the representations of the augmented versions of the *i*-th graph ***𝒢***_*i*_. Let,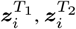 and 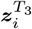 denote the representations of three different sentences describing the *i*-th graph ***𝒢***_*i*_. The objective of contrastive learning is to minimize the contrastive loss for each graph in the mini-batch, considering both cross-modal and self-contrastive losses.

#### 3.3.2 Contrastive Learning Objective

The purpose of the cross-modality contrastive loss is to align representations of molecules and text in the latent space, while the self loss aims to align augmented representations within the same modality.

We define the cross-modality contrastive loss as *ℒ*_*cross*_ and self-contrastive loss as *ℒ*_self_ for the *i*-th graph within a mini-batch of *N* molecular graphs, each with three different augmentations. The final contrastive learning objective combines cross-modal and self-contrastive losses: *ℒ*_MolLM_ = *ℒ*_cross_ + *ℒ*_self_, where 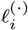 calculates the contrastive loss for the *i*-th graph ***ℒ***_*i*_ considering the respective pairs of multimodal representations.

For an explanation of contrastive loss, *ℒ*_*cross*_, recall that for the *i*-th graph of *N* molecular graphs, 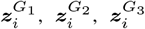, and 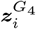 are four representations of augmented graphs and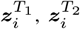, and 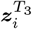 are representations of related text. Equation 6 describes the contrastive loss as a measure of similarity between the graph representations (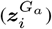) and the corresponding text representations (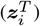) for a given molecular graph. This loss function is formulated as a negative logarithm of the ratio of the exponential similarity between the matching graph and text representations (for a specific graph-text pair) to the sum of exponential similarities across all possible graph-text pairs within the batch of *N* molecular graphs. The term 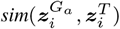 denotes the cosine similarity function between a graph representation and a text representation, and *τ* is a temperature parameter that scales the similarity scores. The loss aims to maximize the numerator, which represents the similarity between matching pairs, and minimize the denominator, which represents the similarity between non-matching pairs. This approach encourages the model to produce graph and text representations that are closer in the embedding space for matching pairs, while pushing non-matching pairs farther apart. Figure 2 displays a t-SNE visualization of molecule embeddings generated by MolLM, illustrating distinct clusterings for each molecule type. This demonstrates the model’s capability to characterize molecules via rich representations.

**Fig. 2.**
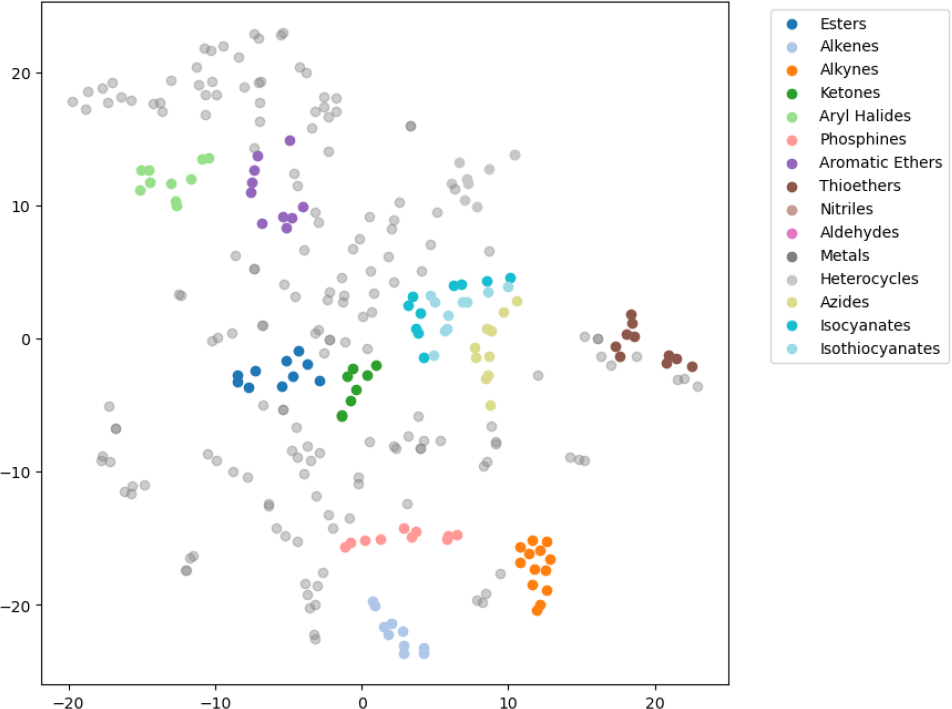
t-SNE visualization of molecule embeddings in MolLM’s joint latent space for select groups of molecules. Colored points represent molecules belonging to the select groups while the gray points represent a variety of other molecules. We displayed 200 data points for a clearer visualization and procured assistance from a chemistry expert to perform annotations of categorization.

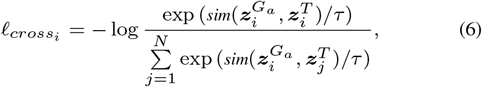

Similarly, for the self loss *ℒ*_self_, recall the same notation for the augmented graphs and related text. This loss function in Equation 7 is defined as the negative logarithm between the ratio of the sum of exponential similarities between all pairs of augmented graph representations (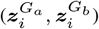) to the sum of all exponential similarities of all such pairs in the graph. The effect of this loss is to ensure that different augmented representations of the same molecular graph are similar to each other in the embedding space. The parameter *τ* is a temperature parameter that scales the similarity scores. This loss encourages the model to push different augmented views of the same graph closer together while pushing those of different graphs farther apart.

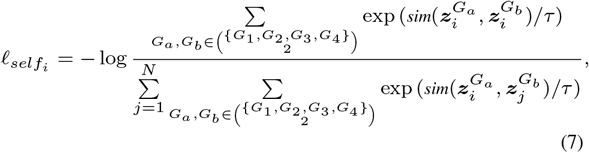

Here, the notation 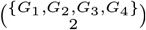 represents the selection of any combination of two different graphs from the set of four. These four graphs represent the molecules modified by four different augmentations.

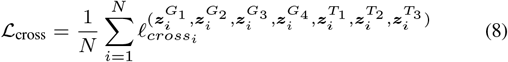

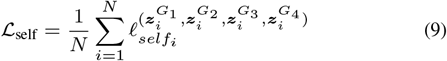

In brief, the contrastive learning objective consists of two components: cross-modal contrastive loss and self-contrastive loss. The cross-modality contrastive loss minimizes the distance between the different modalities (i.e., molecular graphs and related textual descriptions) of the same molecule while maximizing the distance between different molecules. The self-contrastive loss minimizes the distance between different augmentations of the same molecule while maximizing the distance between augmentations of different molecules. Thus, the model learns to generate more robust and semantically meaningful representations for both molecules and text in a joint latent space.

### 3.4 Fine-tuning Downstream Tasks

To showcase the adaptability and practicality of our model across a wide variety of downstream tasks that necessitate both molecular and textual inputs, we fine-tune our model on four such tasks. To provide succinct overviews of these tasks, Cross-modality Matching involves retrieving the correct molecular graph or text from a list of data in the opposing modality. Property Prediction is a classification task for various experimental molecular properties, many of which are related to biological activities such as blood-brain barrier penetration (BBBP). Molecule Captioning involves generating a useful text description of a molecule. Molecule Editing involves generating a new molecule given an original molecule and a prompt with a desired property in text form. See Table 2 for a summary of these tasks.

**Table 2.**
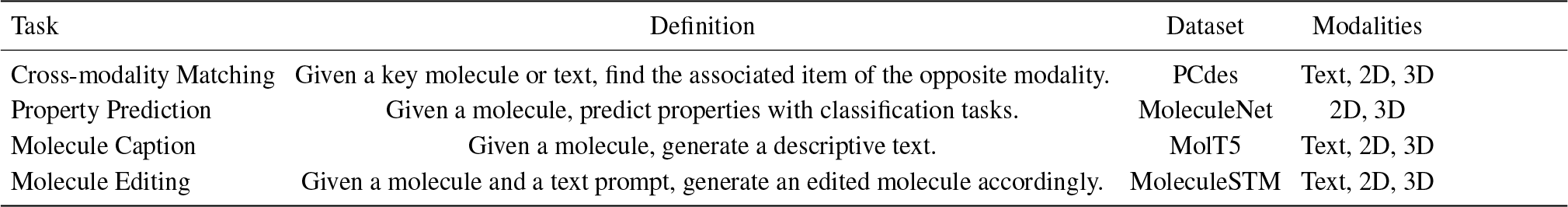
Description of downstream tasks tested along with their associated dataset source (database or paper providing this data) and modalities.

#### 3.4.1 Cross-modality Matching

To implement the Cross-modality Matching task using our model, we begin by using the two encoders to produce embeddings for all of the text and molecules, respectively. A “key” text or molecule is used as a query to find a match in the opposing modality. We generate embeddings for the “key” text or molecule, as well as for the set of texts or molecules that will be used for querying. We then compare the “key” text or molecule against the set of the opposite modality by selecting those with maximal cosine similarity between their embeddings, aiming to find the best match (or matches, in the context of the recall metric). See Figure 3 for a visualization of how the model considers a specific molecule and text pair by computing cosine similarity for the task. For fine-tuning on this task, similar to (Su *et al*., 2022), we split the data into a training set of 10, 500 molecules, a validation set of 1, 500 molecules, and a test set of 3, 000 molecules, while generating their 2D graphs.

**Fig. 3.**
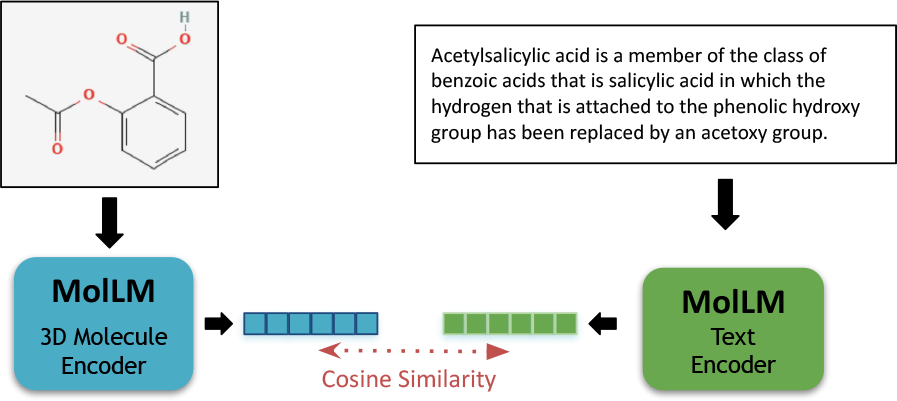
Visualization of the cross-modality matching text for a given pair of molecules and text. Embeddings are generated for each molecule and text using the respective encoders of MolLM. Then, cosine similarity is used to find the most similar pair for top-1 matching, or the 20 most similar for R@20 (Recall), for the matching task.

**Fig. 4.**
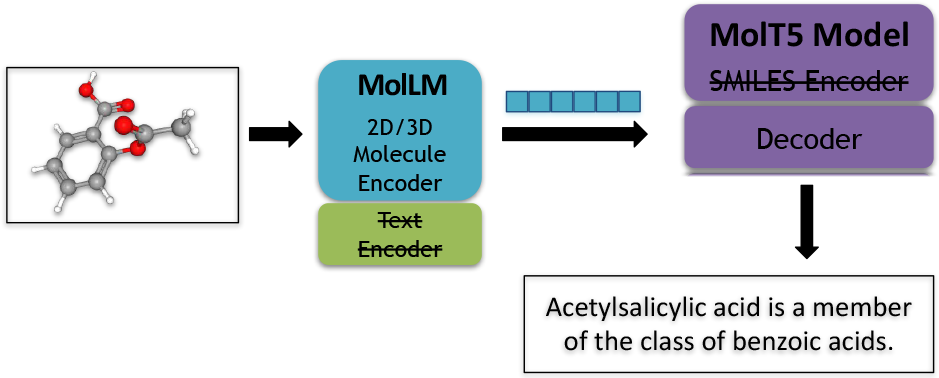
Visualization of the molecule captioning task. We begin with a 2 D/3D molecular graph of acetylsalicylic acid, which is then encoded through MolLM’s molecule encoder. This encoded representation is then decoded through MolT5’s decoder, which produces a molecule-specific caption.

#### 3.4.2 Property Prediction

For the Property Prediction task, which is a classification task, using our model, we employ only the 2D/3D molecular encoder to produce an embedding for the molecule. Then, we attach a graph prediction head to our molecular encoder, which yields a 768-dimensional molecule embedding, as described in Appendix 1.

#### 3.4.3 Molecule Captioning

For the implementation of the Molecule Captioning task, we utilize the 2D/3D molecular encoder of our model to produce an embedding for the molecules, and then forward this embedding through the MolT5l model’s decoder (Edwards *et al*., 2022). We utilize this decoder to generate the descriptions, as our model does not have a decoder for text. We utilize both the small (*∼*77M parameters) and base (*∼* 250M parameters) checkpoints of MolT5. Refer to Figure 6 for a visualization of this technique for generating captions. The -small and -base suffixes on the model names denote the checkpoint of MolT5 used.

#### 3.4.4 Molecule Editing

Finally, for the Molecule Editing task, we have an input molecule and a text prompt. The desired output is a generated molecule that retains similarity to the input molecule while having properties indicated in the text prompt. As MolLM lacks a decoder to directly reverse its embeddings into molecular graphs, we utilize MoFlow (Zang and Wang, 2020), an invertible flow-based molecular model, for generation. We optimize the embedding in MoFlow’s latent space to enable its reversal into the final edited molecule, while computing losses within MolLM’s latent space. Refer to Figure 5 for a diagram of this process. We pass the input molecule and text prompt through the molecule and text encoders of MolLM, respectively, to obtain embeddings of each. In the diagram, these are represented as blue and green rectangles, respectively.

**Fig. 5.**
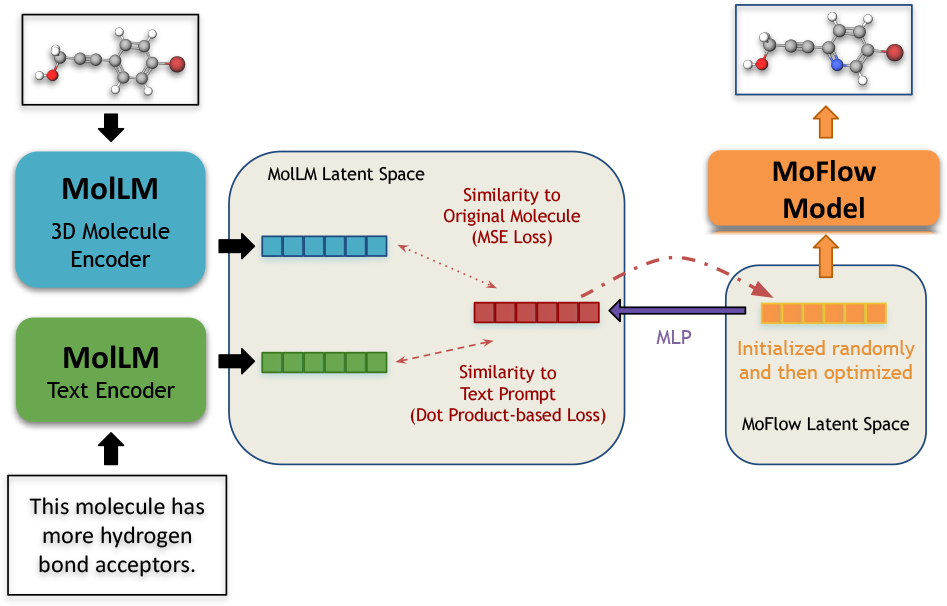
Visualization of molecule editing task. We start with a molecular graph and an instruction on how to edit the molecule. The graph and text are sent to MolLM’s molecule and text encoders, respectively. We optimize an embedding in MoFlow’s latent space but with two losses, to retain similarity to the original molecule while following the text prompt, computed on a translated embedding within MolLM’s latent space. The optimized embedding within MoFlow’s latent space is then reversed through MoFlow to obtain the final molecule.

To begin the optimization process, we initialize an embedding in the MoFlow latent space, represented by the orange rectangle in the diagram, to a random embedding from Gaussian noise. Then, we translate this embedding into an embedding within MolLM’s latent space, in which we denote the intermediate embedding by an MLP, represented by the purple arrow in Figure 5. Refer to Appendix 4 for details of this MLP. Next, to indirectly optimize the embedding within the MoFlow latent space, we compute two losses within the MolLM latent space (as shown in Figure 5 by double-headed red dashed arrows). The first loss aims to maintain similarity to the embedding of the original molecule. The second loss aims to align the embedding of the original molecule with the embedding of the text prompt to achieve the desired molecular properties. See Appendix 3 for details of these losses. After the losses within the MolLM latent space are computed, the gradients with respect to the embedding within the MoFlow latent space, represented by orange in the diagram, are backpropagated, represented by a curved red arrow. Then, these gradients are used to optimize the embedding in MoFlow latent space to minimize both of the aforementioned losses. Thus, the optimized embedding represents a molecule with similarity to the input molecule but possessing the properties described in the text prompt. After 600 optimization steps, we obtain the final state of the embedding. Ultimately, we reverse the embedding, represented by orange in Fig 5, through the invertible MoFlow model to produce the final edited molecule, as illustrated by the orange arrow.

## 4 Results

### 4.1 Cross-modality Matching

**M-T** (molecule-to-text) retrieval begins with a 3D molecular graph and aims to select the most relevant textual descriptions associated with the given molecule. Conversely, **T-M** (text-to-molecule) retrieval starts with a text description and attempts to identify the molecular graph that best corresponds to the description. Both supervised tasks utilize the PCDes dataset (Zeng *et al*., 2022), which is comprised of SMILES strings and their corresponding textual descriptions. The dataset used for zero-shot learning is consistent with the one composed by MoMu. We generate 2D graphs for each example and then convert these SMILES strings into PubChem CIDs.

To incorporate 3D conformation data into our model, we modify each graph by adding 3D information using RDKit’s MMFF94 implementation. Similar to (Su *et al*., 2022), we perform retrieval in randomly selected batches of 64, assessing the average accuracy of top-1 retrieval results and the recall of top-20 (R@20) retrieval results. Specifically, the recall of top-1 retrieval measures how often the model retrieves the correct result as its best match, while the recall of top-20 retrieval measures how often the best match is among what the model deems to be the top 20 most similar results. For each task, we create both sentence-level and paragraph-level tasks. In the sentence-level setting, following (Su *et al*., 2022), we use a randomly chosen sentence from the molecule’s description. In the paragraph-level setting, we utilize the entire description. We adopt Sci-BERT, KV-PLM, and MoMu as baselines.

From Table 3, it is clear that MolLM-3D achieves notable results by outperforming all baselines across each task. This performance is evident in both zero-shot and supervised settings, encompassing sentence-level and paragraph-level tasks. Particularly impressive is the M-T R@20 (Recall) score of **92.05±1.03** in the supervised sentence-level task for MolLM-3D, indicating a high recall rate and suggesting that the model is highly effective in retrieving relevant textual descriptions for given molecular structures. Similarly, T-M accuracy and recall at both the sentence level and paragraph level in the supervised setting show a marked improvement over the other models, with MolLM-3D scoring **66.11** and **91.75**, respectively. As MolLM-3D consistently outperforms MolLM-2D, these results demonstrate the efficacy of incorporating 3D structural information into the retrieval process.

**Table 3.**
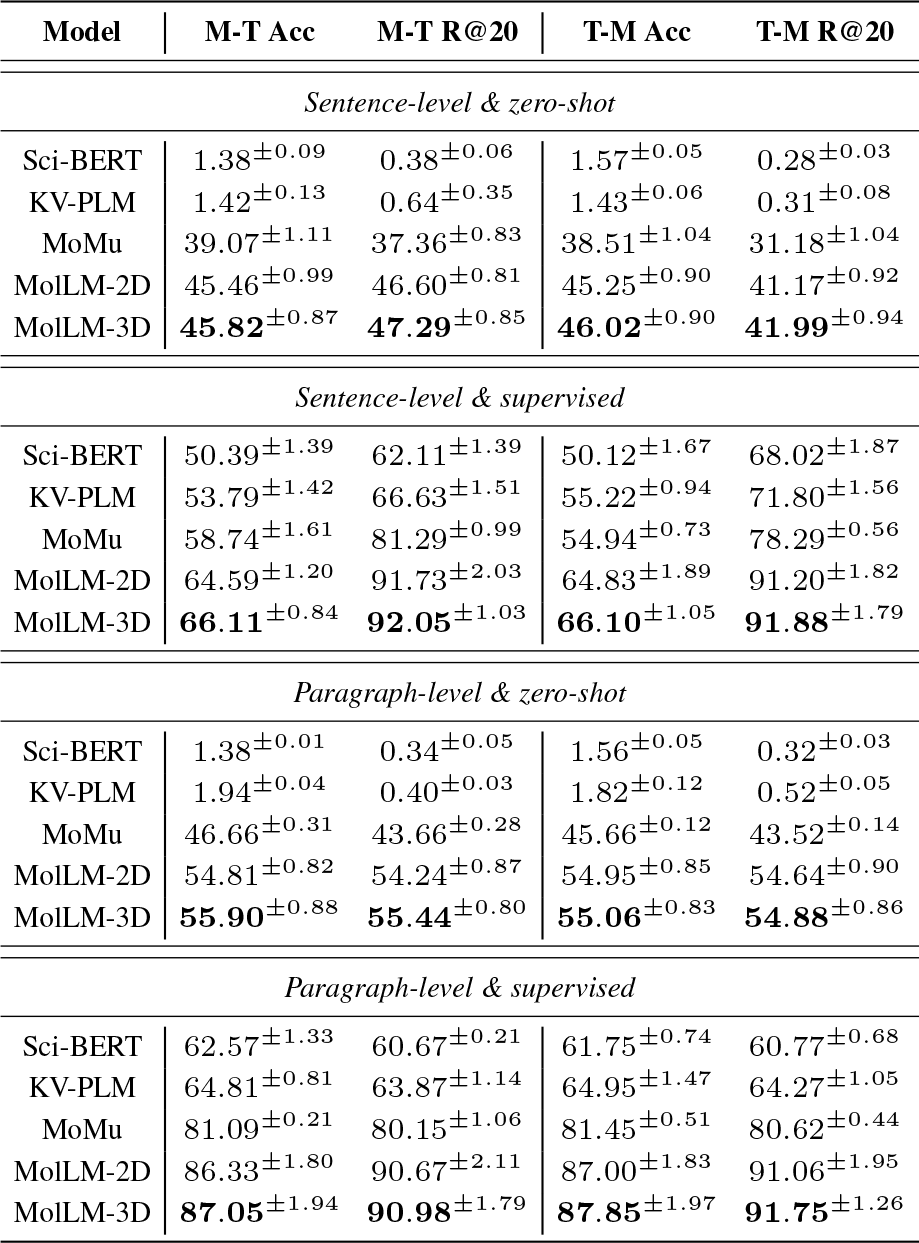
Molecule-to-text (M-T) and text-to-molecule (T-M) retrieval results, presented across four different settings: sentence-level zero-shot prediction, sentence-level supervised fine-tuning, paragraph-level zero-shot prediction, and paragraph-level supervised fine-tuning. R@20: top 20 recall.

### 4.2 Property Prediction

Property prediction is a downstream task widely used in prior work, such as (Su *et al*., 2022) and (Liu *et al*., 2023b). Similar to these studies, we utilize the MoleculeNet (Wu *et al*., 2017b) benchmark, aimed at assessing how effectively a pre-trained molecular graph encoder can adapt to various disparate classification tasks. For example, the TOX21 and HIV datasets might prompt a model to classify new molecular graphs for toxicity markers or anti-HIV activity, respectively. We directly compare our model with relevant approaches such as MoMu and KV-PLM, which use a similar multimodal strategy to our model. As shown in Table 4, our model excels in property prediction tasks compared to the relevant baselines. In zero-shot scenarios, MolLM scores **72.4** while MoMu scores **66.96** on average, MolLM outperforms MoMu by up to **8.14%**, demonstrating enhanced generalization capabilities. Upon fine-tuning, MolLM’s performance surpasses that of the baselines by significant margins, evident in datasets such as TOX21, where it increases from **75.6%** to **80.0%**, and BACE, where it improves from **77.1%** to **84.1%**. These enhancements underscore the efficacy of our expressive molecular encoder, which captures important 3D information. This aspect becomes crucial for classification tasks where considering molecular structure is important.

**Table 4.**
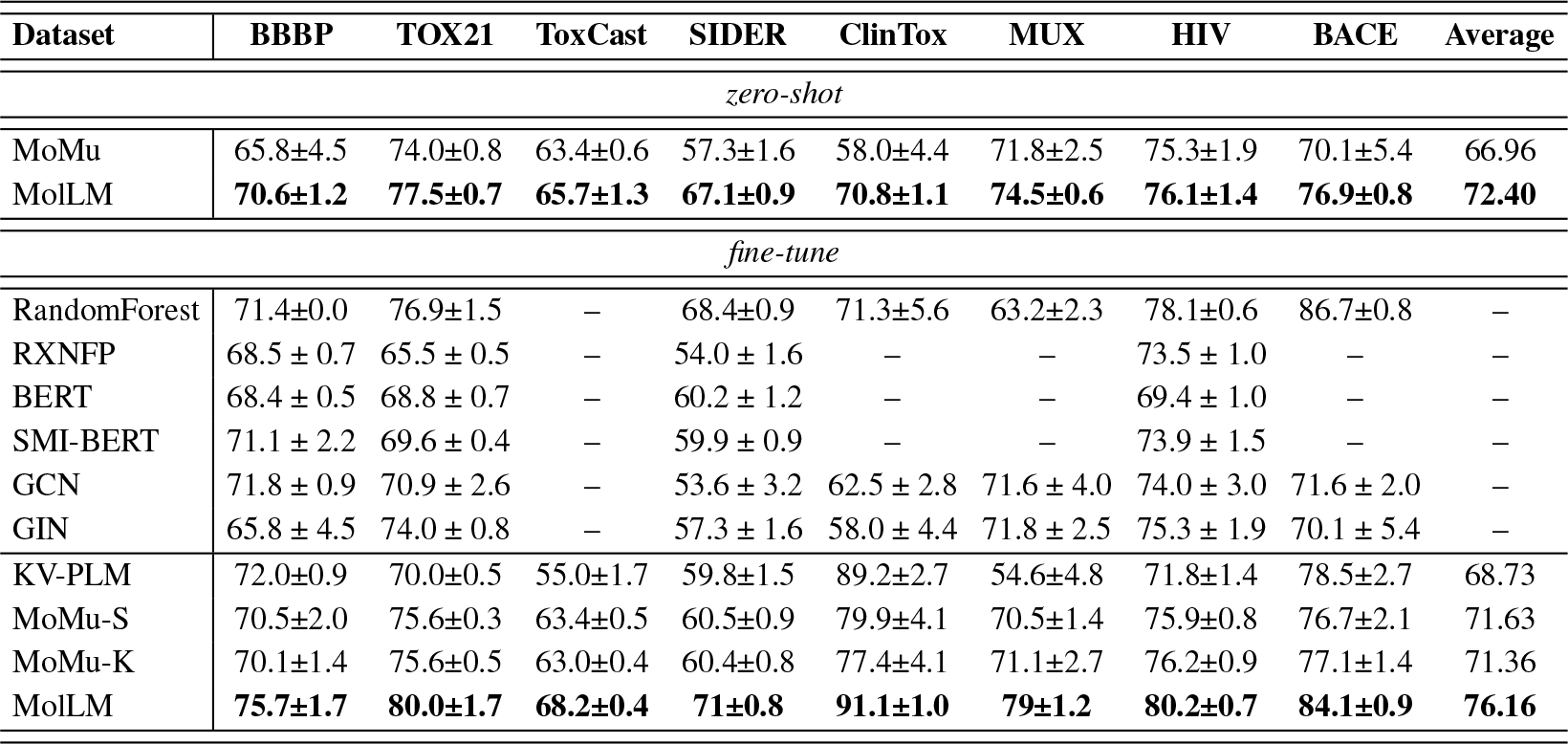
Results of the property prediction task compared with other relevant baselines that also utilize a multimodal biomedical text and molecule model. We also include other fine-tuned models for comparison, but only directly compare against the aforementioned relevant baselines.

| Dataset | BBBP | TOX21 | ToxCast | SIDER | ClinTox | MUX | HIV | BACE | Average |
| --- | --- | --- | --- | --- | --- | --- | --- | --- | --- |
| *zero-shot* |  |  |  |  |  |  |  |  |  |
| MoMu | 65.8±4.5 | 74.0±0.8 | 63.4±0.6 | 57.3±1.6 | 58.0±4.4 | 71.8±2.5 | 75.3±1.9 | 70.1±5.4 | 66.96 |
| MolLM | **70.6±1.2** | **77.5±0.7** | **65.7±1.3** | **67.1±0.9** | **70.8±1.1** | **74.5±0.6** | **76.1±1.4** | **76.9±0.8** | **72.40** |
| *fine-tune* |  |  |  |  |  |  |  |  |  |
| RandomForest | 71.4±0.0 | 76.9±1.5 | — | 68.4±0.9 | 71.3±5.6 | 63.2±2.3 | 78.1±0.6 | 86.7±0.8 | — |
| RXNFP | 68.5 ± 0.7 | 65.5 ± 0.5 | — | 54.0 ± 1.6 | — | — | 73.5 ± 1.0 | — | — |
| BERT | 68.4 ± 0.5 | 68.8 ± 0.7 | — | 60.2 ± 1.2 | — | — | 69.4 ± 1.0 | — | — |
| SMI-BERT | 71.1 ± 2.2 | 69.6 ± 0.4 | — | 59.9 ± 0.9 | — | — | 73.9 ± 1.5 | — | — |
| GCN | 71.8 ± 0.9 | 70.9 ± 2.6 | — | 53.6 ± 3.2 | 62.5 ± 2.8 | 71.6 ± 4.0 | 74.0 ± 3.0 | 71.6 ± 2.0 | — |
| GIN | 65.8 ± 4.5 | 74.0 ± 0.8 | — | 57.3 ± 1.6 | 58.0 ± 4.4 | 71.8 ± 2.5 | 75.3 ± 1.9 | 70.1 ± 5.4 | — |
| KV-PLM | 72.0±0.9 | 70.0±0.5 | 55.0±1.7 | 59.8±1.5 | 89.2±2.7 | 54.6±4.8 | 71.8±1.4 | 78.5±2.7 | 68.73 |
| MoMu-S | 70.5±2.0 | 75.6±0.3 | 63.4±0.5 | 60.5±0.9 | 79.9±4.1 | 70.5±1.4 | 75.9±0.8 | 76.7±2.1 | 71.63 |
| MoMu-K | 70.1±1.4 | 75.6±0.5 | 63.0±0.4 | 60.4±0.8 | 77.4±4.1 | 71.1±2.7 | 76.2±0.9 | 77.1±1.4 | 71.36 |
| MolLM | **75.7±1.7** | **80.0±1.7** | **68.2±0.4** | **71±0.8** | **91.1±1.0** | **79±1.2** | **80.2±0.7** | **84.1±0.9** | **76.16** |

**Table 5.** Results of molecule editing task presented on different prompts. The satisfactory hit ratio results for our model’s performance on this task for the selected 200 molecules are scaled to a range of 0.0-100.0.

| Model | Prompt | Hit Ratio |
| --- | --- | --- |
| MolLM-3D | This molecule has more hydrogen bond acceptors | 46.0 |
| MolLM-3D | This molecule has more hydrogen bond donors. | 54.5 |
| MolLM-3D | This molecule is like a drug. | 5.00 |
| MolLM-3D | This molecule is not like a drug. | 81.5 |
| MolLM-3D | This molecule has high permeability. | 33.5 |
| MolLM-3D | This molecule has low permeability. | 56.5 |
| MolLM-3D | This molecule is soluble in water. | 76.5 |
| MolLM-3D | This molecule is insoluble in water. | 21.5 |

### 4.3 Molecule Captioning

We utilize the molecule captioning task as outlined in (Edwards *et al*., 2022). In this task, our model generates text relating to a molecule based on its SMILES string, 2D graphs, and 3D graphs. For baseline comparisons, we select MolT5 (Edwards *et al*., 2022), which generates captions by inputting SMILES strings into their encoder-decoder transformer architecture. Another baseline we use is MoMu (Su *et al*., 2022), which expands upon this process by appending 2D structural data to the SMILES embedding inputs. To improve our model’s understanding of the molecular structure and generate more accurate descriptions, we integrate 3D data into the input. Using the CheBI-20 dataset as in (Edwards *et al*., 2022), we compare the performance of MolLM, MoMu, and MolT5 using four different captioning metrics, as shown in Fig 6. In summary, MolLM performs equivalently or slightly better than each baseline. What this reveals is that, even without pre-training a text decoder, MolLM representations obtained through the robust encoder are sufficient in generating textual descriptions of molecules. This outcome underscores the versatility of our approach, its ready plug-and-play compatibility, and its ability to conserve computational resources. It can effortlessly leverage other model’s decoder to meet varying needs. Simultaneously, we acknowledge that this dataset has its limitations. Its small scale may not fully demonstrate the potential of our model.

**Fig. 6.**
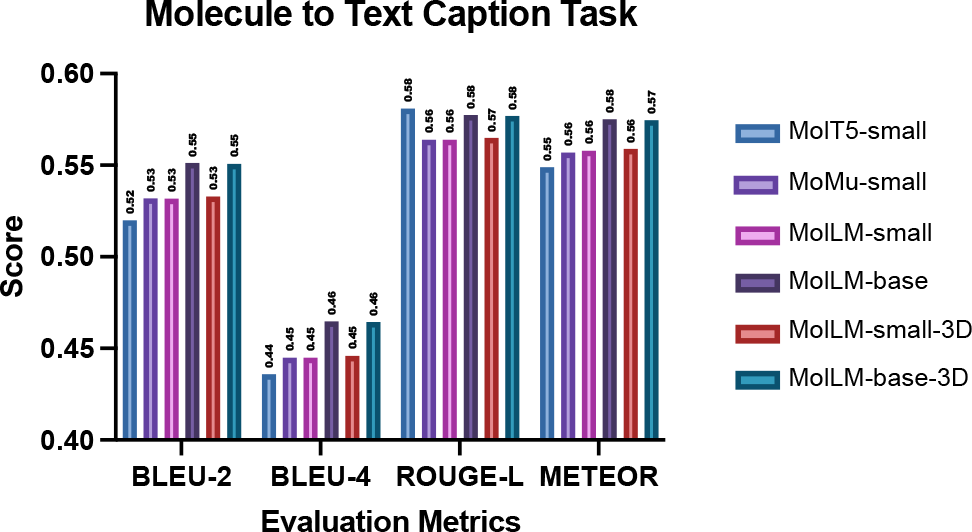
Our results for the molecule captioning task. We utilize the ChEBI dataset (Edwards et al., 2022) with 33,010 molecules. We use BLEU-2, BLEU-4, ROUGE-L, and METEOR to compare performance with MolT5 and MoMu as baselines. “-small” & “-base” suffixes on model names indicate the size of MolT5 decoder because MolLM is encoder-only.

**Fig. 7.**
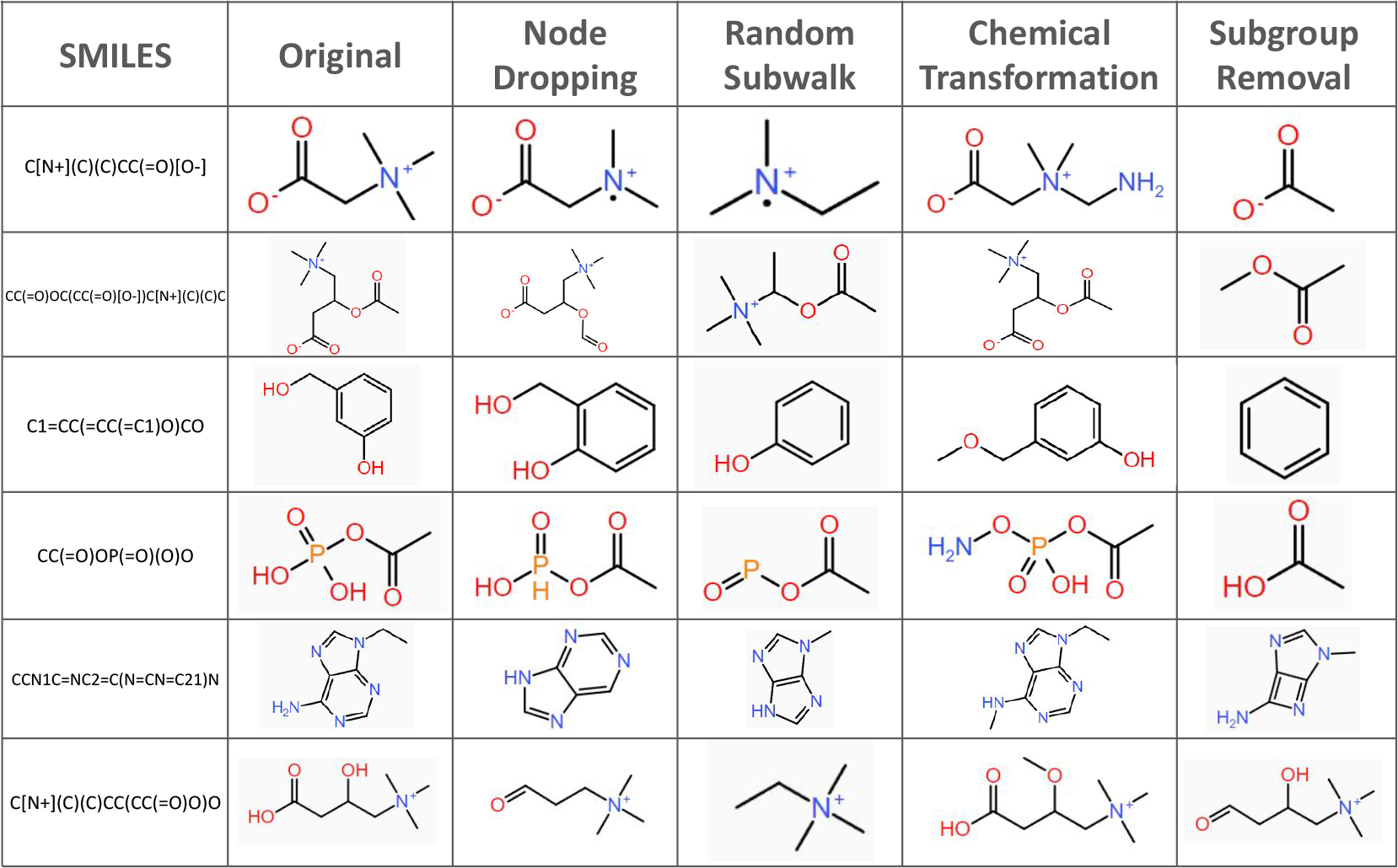
Examples of different molecule augmentations. For each molecule, their SMILES string, original molecular graph, and the molecular graph of each of the four augmentations are shown.

### 4.4 Molecule Editing

Following the procedure and settings outlined in (Liu *et al*., 2023b), we begin with 200 randomly sampled molecules from ZINC (Sterling and Irwin, 2015) along with a brief text editing prompt as inputs. The task is to perform single-objective editing with prompts, provided by Liu *et al*., such as “this molecule has high permeability.” These prompts direct the model to edit a molecule toward a specific molecular property. We evaluate each of these metrics by their satisfactory “hit” ratio, which measures the ratio of the 200 molecules that the model can modify following the prompt, where a successful modification is defined as a “hit.” The determination of hits relies on calculated molecular metrics, each corresponding to a prompt. For example, a satisfactory hit for the “high permeability” prompt would involve the model outputting a molecule with increased permeability, measured by a lower calculated topological polar surface area. For other properties—hydrogen bond acceptors, hydrogen bond donors, drug-likeness, and solubility—the respective calculated properties used to determine hits are calculated hydrogen bond acceptors, calculated hydrogen bond donors, QED drug-likeness (Bickerton *et al*., 2012), and Wildman-Crippen LogP values (Wildman and Crippen, 1999).

Upon examining the results for molecule editing, we noted distinct hit ratios for diverse properties. The model achieved a success ratio of 46.0% for increasing hydrogen bond acceptors and 54.5% for bolstering hydrogen bond donors. Yet, there was a clear dichotomy when modifying drug-likeness - the model attained a low hit ratio of 5.0% for rendering molecules more drug-like, compared to a considerably higher ratio of 81.5% for achieving the opposite. This discrepancy stems from the stark differences in the sizes of these two subtasks’ respective search spaces. In terms of editing for permeability, MolLM increases permeability with a success ratio of 33.5% and reduces it with an efficacy of 56.5%. This disparity could be possibly linked to the artificially set parameters used to define a satisfactory hit. Interestingly, the model performed well in enhancing solubility in water, achieving a hit ratio of 76.5%. In sum, these results suggest that while MolLM shows potential in certain molecular editing tasks, there is still scope for refinement to bolster. The nature of the molecule editing task prevents us from making a direct comparison with MoleculeSTM, as the input molecules vary.

## 5 Discussion

We selected a wide range of tasks to showcase MolLM’s flexibility in multiple contexts. For **property prediction**, our model does not explicitly use the text encoder. However, even without direct usage of text, the embeddings generated through the molecule encoding are still influenced by the contrastive learning process, thereby indirectly benefiting from the biomedical text knowledge. This concept is evident in our utilization of fixed encoders for tasks, such as employing MolT5 for molecule captioning and incorporating MoFlow for molecule generation. In these tasks, we maintain our encoder in a fixed state during fine-tuning, which enables a better understanding of our fine-tuning effects, prevents overfitting, and reduces computational costs. These tasks can perform well because the encoders are pre-trained and our multimodal approach provides sufficient context before fine-tuning.

Our results significantly surpassed current state-of-the-art baselines in **cross-modality matching** and **property prediction** tasks. In the matching task, incorporating 3D data yielded substantially better results compared to its 2D counterpart, with both outperforming all baselines. These outcomes not only underscore the importance of the inclusion of multimodal contexts for pre-training but also highlight the particular significance of 3D encoding.

In **molecule captioning**, our model also achieved competitive results compared to baselines. We attribute this success primarily to the size of our dataset, which is larger than other text-based datasets. Interestingly, we observed that the inclusion of 3D data does not significantly improve results in this task. This discrepancy could be due to the heavily text-focused nature of the task, where the text may implicitly not require a large amount of molecule structure data. In other words, the captions themselves may not strongly correlate with molecular structure data. Conversely, captions such as “the molecule is symmetrical” may be more likely to benefit from incorporating molecular structural data.

In **molecule editing**, the performance varied significantly depending on the editing task. This variation could be due to the difficulty of specific prompts, as some prompts yielded successful hits for a much narrower range of similar molecules to the original. However, our model has constraints limiting the extent of molecule editing to avoid excessive alterations. For example, in the case of the solubility prompt, over-editing could potentially result in a highly soluble molecule that bears no resemblance to the original. Additionally, the variation in performance among the prompts might stem from inherent limitations within the training data, particularly in how frequently the text related to a specific prompt occurs.

Upon further investigation, we found that many edits resulted in molecules vastly different from the original, suggesting that a high hit ratio might also arise from overly editing a molecule. This highlights issues with the metric itself, suggesting a potential need for developing a more robust metric for evaluation. The existing metric solely considers changes in relevant calculated properties while ignoring the similarity to the original molecule. A key aspect of molecule editing is retaining high similarity to the original, yet the metric encourages edits that strongly favor the prompt without considering the original structure. A better evaluation metric could involve a combination of both similarity to the original molecule and adherence to the prompt.

The value of integrating 3D information cannot be overstated. Through a multimodal approach, we capture a richer representation of molecules, highlighting the interplay between their structural and spatial features. Additionally, the quality and size of the utilized datasets play a pivotal role in the performance of these models. Based on the demonstrated effectiveness of this architecture, molecular models should progress towards embracing a multimodal approach that incorporates 3D data.

## 6 Conclusion

MolLM, with its emphasis on including 3D positional data, exhibits robust performance across a variety of downstream tasks including cross-modality matching, property prediction, captioning, and editing. This transformative approach underscores the importance of incorporating a richer molecular representation, highlighting the nuanced interplay between structural and spatial features. Additionally, it emphasizes the important role of incorporating biomedical knowledge via textual inputs. In the future, this textual input could be further augmented using GPT to enrich the available data pool for pre-training. Our model demonstrates a promising future for utilizing computational methods that incorporate the wealth of existing biomedical literature to advance complex tasks such as drug discovery that require a complex understanding that transcends multiple modalities.

## 1 Property Prediction Head

Below are the details of the prediction head we append to the base molecular encoder within our model to adapt to the property prediction task.

The main motivations behind the prediction head layer involve reducing features to learn the most important features for property prediction in the linear layers, ensuring a uniform distribution of outputs with the normalization layers for training stability, and considering dropout to avoid both underfitting and overfitting (Liu *et al*., 2023d). We perform a grid search to obtain the final prediction head layer dimensions and hyperparameters.

- Linear layer: 768 *×* 512
- 1D Batch Normalization: 512 features
- Dropout: rate of 0.2
- Activation: ReLU
- Linear layer: 512 *×* 300
- Activation: ReLU
- Dropout: rate of 0.2
- Linear layer: 300 *× C*, where *C* represents the number of categories for the specific property prediction task.

## 2 Use of RDKit

We use RDKit to represent molecules from SMILES strings and obtain their graph structure prior to transforming their structures to our specific graph format with PyTorch Geometric. The motivation behind having an upper threshold of RMSD after performing molecular augmentation is to ensure that the augmentation does not result in a molecule that is too semantically different for use in contrastive learning.

## 3 Molecule Augmentation Examples

We utilize four different molecule augmentations during our pre-training process: Node Dropping, Random Subwalk, Chemical Transformation, and Subgroup Removal. We chose these augmentations as they edit the molecules in a manner in which we believe the semantic meaning of the molecule is kept but is still augmented enough for the model to be more robust and explore the latent space. A more detailed explanation of each of these processes can be found in section 3.3.1. Figure 7 depicts five examples of how our augmentations would take an original molecule and convert it into their augmented versions. Let’s look at a few examples. The first row shows the molecule Betaine. We can see the effect of node dropping, as one of the carbons has been dropped from the nitrogen. The random subwalk has started on the carbon atom, which was connected to the oxygen atoms, and walked through the graph. The chemical transformation has added an amine group to one of the carbon atoms. The subgroup removal has left only one subgroup of the Betaine, based on BRICS decomposition. Another molecule we can examine is in the fourth row: acetylphosphate. We can see that node dropping has caused an oxygen to be dropped from the phosphorus atom. The random subwalk has taken a walk of the molecular graph, starting from the oxygen atom that is double-bonded to the phosphorus atom. The chemical transformation has added an amine group to an oxygen atom. The subgraph removal has left us with a random subgroup of the graph, as dictated by BRICS decomposition.

## 4 Training Implementation Details

### 4.1 Cross-modality Matching

We utilize a contrastive loss between the graph representations and text representations for fine-tuning. See Equation 10, where *m* is the margin to control how far apart embeddings of negative pairs should be relative to positive pairs and *<*_*ij*_ is the Kronecker delta function.

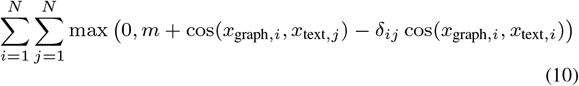

Specifically, we utilize *m* = 0.2 throughout all of our fine-tuning of this task. We fine-tune for 60 epochs with a batch size of 64 at a learning rate of 5*·*10^*-*5^ for each subtask.

### 4.2 Property Prediction

For the fine-tuned tasks, which are all classification tasks, we utilize a Binary Cross-Entropy loss. See Equation 11 where *N* is the number of samples, *ŷ*_*i*_ is the predicted value, and *y*_*i*_ is the target label.

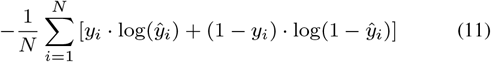

We fine-tune for 200 epochs with a batch size of 32 with a learning rate of 10^*-*4^. See Appendix 1 for details on the linear prediction head for the classification task.

### 4.3 Molecule Caption

We utilize a Cross-Entropy Loss between the predicted captions and target captions. See Equation 12 for details on this loss, where *p*(*y*_*ij*_ |*x*_*i*_) is the model’s probability for a token given the input concatenated SMILES string embedding and molecule embedding, *x*_*i*_.

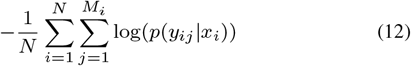

We fine-tune for 10 epochs with a batch size of 16 with a learning rate of 10^*-*4^.

### 4.4 Molecule Editing

Finally, for the molecule editing task, we do not fine-tune our model as the task involves *de novo* generation. Instead, we optimize the predicted molecule such that its embedding through MolLM is more aligned with the text prompt’s embedding within the MolLM latent space. We utilize a mean squared error (MSE) between the predicted molecule embedding and the original molecule embedding to maintain similarity to the original molecule. See Equation 13 for the MSE loss, where *z*_pred_ is the predicted molecule embedding, *z*_orig_ is the original molecule embedding, and *N* is the number of samples.

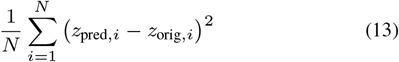

Additionally, we incorporate a Dot Product-based loss to ensure alignment between the predicted molecule’s embedding and the text prompt’s embedding. See Equation 14 for the Dot Product-based loss, where *z*_*text*_ is the text prompt’s embedding.

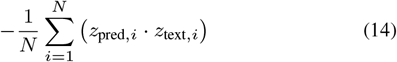

This dual approach with a linear combination of these two losses for similarity retention and alignment to the prompt ensures that the edited molecules are related to the original while being edited in a manner relevant to the prompt.

## 5 Embedding Translation MLP

The weights of this MLP for projection from MoFlow latent space to MolLM latent space were trained to minimize cosine similarity between the translated embeddings and the actual MolLM embedding for molecules within the ZINC (Sterling and Irwin, 2015) dataset. The cosine similarity loss for *N* pairs, where *z*_*pred*_ is the predicted translation and *z*_*true*_ is the true embedding through MolLM’s molecule encoder, is

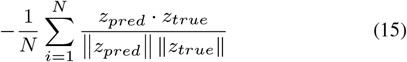

## 6 Model Architecture Explanation

Our choice of model architecture was based on our selection of molecular representations and the advantages offered by Transformers in encoding molecules. For molecular representation, we examined prior works to inform our approach. While SMILES and SELFIES strings are simple and directly leverage natural language pipelines, previous literature has highlighted limitations in these representations. For instance, SMILES strings can represent the same molecule in multiple ways, leading to unnecessary data redundancy. Additionally, both SMILES and SELFIES representations lack the ability to encode 2D/3D spatial information, leading to decreased performance in certain tasks.

Thus, we chose to focus on graph-based molecular representations. Our reasoning for continuing to use Transformers to encode molecules stems from the architecture’s superior capability to handle sequences. In contrast to traditional GNN or GIN models, which primarily emphasize the graph structure of molecules, Transformers allow for more flexible encoding. The attention mechanism within Transformers captures molecular patterns, while the encoder-decoder structure allows us to easily leverage multimodal inputs for pre-training. Consequently, we can effectively utilize textual data as inputs, enabling our model to understand domain-specific knowledge associated with a given molecule.

## 7 Future Direction

In terms of future directions, there is an evident need for more extensive and better-curated datasets. Having larger datasets with a stronger correlation between text and molecular representations would enable the model to establish clearer distinctions among molecules. Automatically identifying such text in large text corpora poses challenges because the mention of molecules in certain texts does not guarantee their relevance. Thus, an improvement in the current text sampling method is necessary. For example, future studies could leverage language models to sift through academic texts or include additional quantitative data from other sources. Exploring improvements in downstream tasks is also important.

Introducing novel downstream tasks could allow us to validate the model’s applicability across a broader spectrum of challenges in molecular biology and chemistry. Additionally, there exist numerous potential directions for improving the downstream tasks utilized by MolLM. For example, a major weakness of the molecule generation task lies in the fact that it sometimes generates infeasible molecules. Future works could aim to validate these generated molecules for chemical validity, chemical stability, or feasibility of synthesis. This could be achieved by employing techniques such as Reinforcement Learning from Human Feedback (Ziegler *et al*., 2019; Stiennon *et al*., 2020), creating a reward function based on expert human knowledge to fine-tune the generation model. Leveraging transfer learning from models like GPT-4 could also aid in this improvement.

For our molecule editing task, future work could explore evaluating how well MoleculeSTM, our model, and other similar models allow for dynamic adjustment of the weighting between emphasizing retaining original molecular features and chemical changes that favor the given prompt. The motivation behind this is that there is likely often the case where a molecule already has many desirable properties and minimal editing is wanted to achieve a prompt. Moreover, a better metric could even consider the ratio of change in the molecule in favor of the prompt to how much the molecule is chemically altered. Our experiments also shed light on the lack of reliable baselines, especially in *de novo* generation, where there is no gold standard. This poses questions about the veracity and stability of generated molecules and presents the opportunity for the creation and adoption of better baselines.

Additionally, the editing task highlights the usefulness of incorporating natural language prompts as input into the model for powerful biomedical-related tasks. Exploring methods to enhance the robustness of this editing task, experimenting with various prompts, and proposing new tasks that involve natural language prompts could lead to more powerful tools leveraging the utility and ease of using natural language prompts.

Furthermore, the pre-training technique could be expanded upon in a manner more advanced than just curating larger or higher-quality datasets. There is also an avenue to explore utilizing much larger language models, such as ChatGPT (Jahan *et al*., 2023) or LLaMA, as an agent for biomedical tasks.

Finally, exploring different molecular encoding methods could yield promising results. For example, while Transformer-M combines 2D/3D data linearly, future works could experiment with non-linear combinations of 2D and 3D data. This approach could enhance expressiveness, enabling the model to better discern the nuances between 2D and 3D representations.

## References

An, X. et al. (2022). Representation of molecules for drug response prediction. Briefings in Bioinformatics, 23(1).

Bickerton, G. R. et al. (2012). Quantifying the chemical beauty of drugs. Nature Chemistry, 4(2), 90–98.

Chen, J. and Zhang, L. (2021). A survey and systematic assessment of computational methods for drug response prediction. Briefings in bioinformatics, 22(1), 232–246.

Chilingaryan, G. et al. (2022). Bartsmiles: Generative masked language models for molecular representations.

Coley, C. W. et al. (2017). Convolutional embedding of attributed molecular graphs for physical property prediction. Journal of chemical information and modeling, 57(8), 1757–1772.

Devinyak, O. et al. (2014). 3d-morse descriptors explained. Journal of Molecular Graphics and Modelling, 54, 194–203.

Devlin, J. et al. (2019). Bert: Pre-training of deep bidirectional transformers for language understanding. In Proceedings of the 2019 Conference of the North American Chapter of the Association for Computational Linguistics: Human Language Technologies.

Edwards, C. et al. (2022). Translation between molecules and natural language. In Y. Goldberg, Z. Kozareva, and Y. Zhang, editors, Proceedings of the 2022 Conference on Empirical Methods in Natural Language Processing, pages 375–413, Abu Dhabi, United Arab Emirates. Association for Computational Linguistics.

Gu, Y. et al. (2021). Domain-specific language model pretraining for biomedical natural language processing. ACM Transactions on Computing for Healthcare (HEALTH), 3(1), 1–23.

Hu, W. et al. (2020). Open graph benchmark: Datasets for machine learning on graphs. Advances in neural information processing systems, 33, 22118–22133.

Hwang, D. et al. (2020). Comprehensive study on molecular supervised learning with graph neural networks. Journal of Chemical Information and Modeling, 60(12), 5936–5945.

Jiang, J. et al. (2021). Ggl-tox: geometric graph learning for toxicity prediction. Journal of chemical information and modeling, 61(4).

Kajita, S. et al. (2017). A universal 3d voxel descriptor for solid-state material informatics with deep convolutional neural networks. Scientific reports, 7(1), 16991.

Kim, S. et al. (2023). Pubchem 2023 update. Nucleic acids research doi:10.1093/nar/gkac956.

Kuenzi, B. M. et al. (2020). Predicting drug response and synergy using a deep learning model of human cancer cells. Cancer cell, 38(5), 672–684.

Kuzminykh, D. et al. (2018). 3d molecular representations based on the wave transform for convolutional neural networks. Molecular pharmaceutics, 15(10), 4378–4385.

Landrum, G. (2023). Rdkit: Open-source cheminformatics.

Li, S. et al. (2022a). Geomgcl: Geometric graph contrastive learning for molecular property prediction. In Proceedings of the AAAI conference on artificial intelligence, volume 36, pages 4541–4549.

Li, Z. et al. (2022b). Deep learning methods for molecular representation and property prediction. Drug Discovery Today, page 103373.

Liu, J. et al. (2023a). The prediction of molecular toxicity based on bigru and graphsage. Computers in Biology and Medicine, 153, 106524.

Liu, S. et al. (2022). Pre-training molecular graph representation with 3d geometry. In International Conference on Learning Representations.

Liu, S. et al. (2023b). Multi-modal molecule structure-text model for text-based retrieval and editing. Nature Machine Intelligence, 5(12), 1447–1457.

Liu, Y. et al. (2023c). Molrope-bert: An enhanced molecular representation with rotary position embedding for molecular property prediction. Journal of Molecular Graphics and Modelling, 118, 108344.

Lo, K. et al. (2020). S2ORC: The semantic scholar open research corpus. In Proceedings of the 58th Annual Meeting of the Association for Computational Linguistics, pages 4969–4983, Online. Association for Computational Linguistics.

Luo, R. et al. (2022a). Biogpt: generative pre-trained transformer for biomedical text generation and mining. Briefings in Bioinformatics, 23(6), bbac409.

Luo, S. et al. (2022b). One transformer can understand both 2d & 3d molecular data. arXiv preprint arXiv:2210.01765.

Miao, Y. et al. (2023). Recent advances in toxicity prediction: Applications of deep graph learning. Chemical Research in Toxicology, 36(8), 1206–1226.

Pogány, P. et al. (2018). De novo molecule design by translating from reduced graphs to smiles. Journal of chemical information and modeling, 59(3), 1136–1146.

Radford, A. et al. (2018). Improving language understanding by generative pre-training. OpenAI blog.

Raffel, C. et al. (2020). Exploring the limits of transfer learning with a unified text-to-text transformer. Journal of Machine Learning Research, 21, 1–67.

Rong, Y. et al. (2020). Self-supervised graph transformer on large-scale molecular data. Advances in Neural Information Processing Systems, 33, 12559–12571.

Ross, J. et al. (2022). Large-scale chemical language representations capture molecular structure and properties. Nature Machine Intelligence, 4(12), 1256–1264.

Scholkopf, B. et al. (1997). Comparing support vector machines with gaussian kernels to radial basis function classifiers. IEEE Transactions on Signal Processing, 45(11), 2758–2765.

Singhal, K. et al. (2023). Large language models encode clinical knowledge. Nature, pages 1–9.

Stärk, H. et al. (2022). 3d infomax improves gnns for molecular property prediction. In International Conference on Machine Learning, pages 20479–20502. PMLR.

Sterling, T. and Irwin, J. J. (2015). Zinc 15 – ligand discovery for everyone. Journal of Chemical Information and Modeling, 55(11), 2324–2337.

Su, B. et al. (2022). A molecular multimodal foundation model associating molecule graphs with natural language. arXiv preprint arXiv:2209.05481.

Thomas, N. et al. (2018). Tensor field networks: Rotation-and translation-equivariant neural networks for 3d point clouds. arXiv preprint arXiv:1802.08219.

Vaswani, A. et al. (2017). Attention is all you need. Advances in neural information processing systems, 30.

Wang, S. et al. (2019). Smiles-bert: Large scale unsupervised pre-training for molecular property prediction. In Proceedings of the 10th ACM International Conference on Bioinformatics, Computational Biology and Health Informatics, BCB ‘19, page 429–436, New York, NY, USA. Association for Computing Machinery.

Wang, Y. et al. (2022a). Improving molecular contrastive learning via faulty negative mitigation and decomposed fragment contrast. Journal of Chemical Information and Modeling, 62(11), 2713–2725.

Wang, Y. et al. (2022b). Molecular contrastive learning of representations via graph neural networks. Nature Machine Intelligence.

Wei, J. et al. (2021). Finetuned language models are zero-shot learners. arXiv preprint arXiv:2109.01652.

Weininger, D. (1988). Smiles, a chemical language and information system. 1. introduction to methodology and encoding rules. Journal of chemical information and computer sciences, 28(1), 31–36.

Wen, N. et al. (2022). A fingerprints based molecular property prediction method using the bert model. Journal of Cheminformatics, 14(1), 1–13.

Wildman, S. and Crippen, G. (1999). Prediction of physicochemical parameters by atomic contributions. Journal of Chemical Information and Computer Sciences, 39, 868–873.

Wu, Z. et al. (2017a). Moleculenet: A benchmark for molecular machine learning.

Wu, Z. et al. (2017b). Moleculenet: A benchmark for molecular machine learning.

Xia, J. et al. (2022). A systematic survey of molecular pre-trained models. arXiv preprint arXiv:2210.16484.

Yang, K. et al. (2019). Analyzing learned molecular representations for property prediction. Journal of chemical information and modeling, 59(8), 3370–3388.

Yu, L. et al. (2021). Review of unsupervised pretraining strategies for molecules representation. Briefings in Functional Genomics, 20(5), 323–332.

Zang, C. and Wang, F. (2020). Moflow: an invertible flow model for generating molecular graphs. In Proceedings of the 26th ACM SIGKDD international conference on knowledge discovery & data mining, pages 617–626.

Zeng, Z. et al. (2022). A deep-learning system bridging molecule structure and biomedical text with comprehension comparable to human professionals. Nature communications, 13(1), 862.

Zhang, Z. et al. (2021). Motif-based graph self-supervised learning for molecular property prediction. Advances in Neural Information Processing Systems, 34, 15870–15882.

Zhu, J. et al. (2022). Unified 2d and 3d pre-training of molecular representations.

Zuranski, A. M. et al. (2021). Predicting reaction yields via supervised learning. Accounts of chemical research, 54(8), 1856–1865.

